# Development of specific molecular markers to distinguish and quantify broomrape species in a soil sample from infected field

**DOI:** 10.1101/602284

**Authors:** Radi Aly, Vinay K. Bari, Avishai Londner, Jackline Abu Nassar, Leena Taha-Salaime, Eizenberg Hanan, Ran Lati

## Abstract

Broomrapes (*Orobanche* and *Phelipanche*) are obligate holoparasites that cause heavy damage to numerous crops, reducing the yield and its quality. The parasite develops in the soil and exerts the greatest damage prior to its emergence; therefore the majority of field loss may occur before diagnosis of infection. Because of the parasite tiny seed size (200 to 300 μm) and dormancy for several decades in the field, it is very difficult to diagnose the parasite by conventional methods. Therefore, to restrict the parasite seeds spread and contamination to other commercial fields, development of DNA-based molecular markers to identify and quantify broomrape species in a soil sample is much needed. In this study, we developed a specific molecular marker (RbcL-M) based on *rbcL* (large subunit of the ribulose-bisphosphate carboxylase) gene from *Orobanche crenata* to differentiate between *Orobanche crenata* and *Orobanche cumana.* Likewise, a specific marker (ITS100) based upon unique sequences in the internal transcribed spacer (ITS) regions of the nuclear ribosomal DNA of *Phelipanche aegyptiaca* to quantify three species of the parasite (*P. aegyptiaca, O. crenata* and *O. cumana*) in a soil sample was developed. Genomic DNA was extracted from soil samples artificially infested with broomrape seeds or tissue of *P. aegyptiaca, O. cumana* and *O. crenata* and subjected to PCR analysis. RbcL-M marker successfully amplified a PCR product (1300bp) when *O. crenata* seeds or tissues (collected from several locations in Israel) were added to the soil samples. The same marker amplified a PCR product (1000bp) when *O. cumana* seeds or tissues were added to the soil samples. RbcL-M marker did not amplify soil samples with seeds or tissues of *P. aegyptiaca* or any soil-borne DNA. Furthermore, using ITS-100 marker and Real-Time PCR analysis, allowed quantitative diagnostic of the parasite in a soil sample from infected sunflower field. As expected the universal internal control primer (UCP-555) amplified a PCR product (555bp) when genomic DNA extracted from soil samples with or without broomrape tissues. The development of an efficient, simple and robust molecular marker to detect and distinguish between broomrape species, has a significant insights on assessment the level of infestation and planning eradication program to the parasite in a field crop.

## INTRODUCTION

Broomrapes (*Orobanche* and *Phelipanche* spp.) are member of Orobanchaceae family (C. Parker, 1993), is an underground root parasite mostly affecting agricultural crop plants (McNeal et al., 2013). These parasitic plants are unable to synthesize their photosynthetic assimilate due to lack of chlorophyll; and totally depend on host plants for the germination and growth by feeding through haustorium (Westwood et al., 2010). The Orobanchaceae family contains the largest number of parasitic species that form haustoria in roots, for example *Orobanche, Phelipanche* species (Yoshida et al., 2016). Haustoria are special organ of parasitic plant which invades host tissue and serve as connecting bridge that allows the parasite to obtain water, mineral and nutrient from host (Joel and Losnergoshen, 1994; Westwood et al., 2010).The natural habitats of broomrapes are the warm and temperate regions of the northern hemisphere, including the Mediterranean region (C. Parker, 1993), however branched broomrape *O. ramosa* is native to central and south-western Europe, and considered as a major threat to agricultural crops. They cause extensive damage by reducing the yield of all economically important crops belongs to Solanaceae, Fabaceae, Brasicaceae, Asteraceae and Apiaceae family (Aly, 2007; C. Parker, 1993; Joel, 2006). In Israel the significance of infested fields has increased dramatically during the last decade, causing heavy crop damage or even total yield losses.

In general, parasitized crops suffer from reductions in total biomass at the greatest expense to their productive tissue (Joel, 2006). Broomrapes cause many problems for agriculture farmers because of their tiny seed size (200-300µm) which makes them difficult to detect in harvested crop seeds and in agriculture soil (Joel, 1987). Moreover, small broomrape germinates and grows only in the presence of a susceptible host. In addition, small broomrape seeds can survive more than a decade in the soil (Habimana et al., 2014). Therefore, identification and quantification of broomrape seeds in a soil sample urgently required and will be helpful in prevention of crop damages by these parasites (Joel et al., 1996). The conventional methods of detection and differentiation such as seed coat morphological feature (Joel, 1987), Random amplified polymorphic DNA technique (RAPD) (Katzir et al., 1996; Paran et al., 1997), Intersimple sequence repeat (ISSR) (Benharrat et al., 2002) and high resolution melting analysis (Rolland et al., 2016) has some limitation to distinguish and quantify broomrapes exist in the soil. Recent progress in sequencing broomrape plastid genome, which was made available, providing new insight to develop molecular marker for specific broomrape species identification (Cusimano and Wicke, 2016; Wicke et al., 2013). Genetic or DNA based molecular marker such as internal transcribe spacer (ITS), plastidial ribulose bisphosphate carboxylase (Rubisco) large subunit (*rbcL*) and maturase K (*matK*) are among the most widely used molecular markers in evolutionary, taxonomy, phylogenetic and diversity identification of many organisms including *Orobanche*, have been established in recent years (Agarwal et al., 2008; Park et al., 2008; Schneeweiss et al., 2004).

In order to develop a complementary methodology for mapping parasitic plants in the field, a procedure includes: a known geographical information systems (GIS) for soil sampling that will characterize the spatial variation in the field (Eizenberg et al., 2012), and at the same time, use of molecular markers to identifying broomrapes in a soil sample. Those molecular markers would assist detection and diagnosis of broomrapes species and population level in the soil. In this study we developed a simple and easy method based on *rbcL* gene to differentiate between three broomrape species *O. crenata, O. cumana* and *P. aegyptiaca*, the most common and destructive weed in Israel. We also developed a specific molecular marker to quantify the parasite seeds in soil sample in an infected sunflower field.

## MATERIAL AND METHODS

### Plant material

Seeds, shoots and inflorescences of three broomrape species (*P. aegyptiaca, O. cumana and O. crenata*) were collected from different Israeli locations (Golan heights, Havat Eden-Emic Betshan and Jordan valley respectively). Seeds were harvested from freshly inflorescences and stored at 4°C while shoots and inflorescences were dried and stored at −80°C until used. Two hundred milligrams of parasite plant seeds/tissue were ground manually in presence of liquid nitrogen with the help of pestle and mortar. Total genomic DNA was extracted using the GeneGET Plant genomic DNA extraction kit according to the manufacturer instructions (Thermo Scientific).

### ITS and *rbcL* sequences and primers design

The internal transcribed spacer (ITS) regions of *Phelipanche aegyptiaca* (GenBank-Accession No. AY209327) was used to design ITS100 marker. The PCR product yielded a 100bp fragment in length. UCP-555 was used as universal internal control primer, which amplifies a region of the small subunit of nrDNA (555 bp) from a wide variety of microorganism such as protists, fungi, and plants. The length of the *rbcL* sequences was 1211bp and 1290bp for *O. cumana* (GenBank: AF090349.1) and *O. crenata* (GenBank: AY582191.1) respectively. Sequence similarities between broomrape species were done using Clustal Omega software. Design of primers for the qPCR detection of broomrape species was based on the alignment of the ITS of nuclear ribosomal DNA regions. All primers were designed with Gene Runner software. PCR amplifications of *rbcL* sequences were performed with the RbcL-M-F and RbcL-M-R primers using PCR Ready Mix (BioLabs, Israel), and 50ng DNA-template. The following PCR conditions were used: denaturation for 5 min at 95°C; 32 cycles with 30 second at 95°C, 30 second at 55°C, 1.30 min at 72°C; and final elongation for 10 min at 72°C.

### Soil sampling and extraction of Genomic DNA

Design and mapping the soil samples in the sunflower tested field were performed according to (Eizenberg et al., 2012). Soil samples were collected from the infected field on May and counting the inflorescences was performed on July. Each sample includes 500gm of a soil sample that was introduced into one-litter container, mixed thoroughly with 1.5L of (13.3 M CaCl_2_) solution and kept at room temperature for overnight. The next day, the upper phase including the organic materials and broomrape seeds were collected, dried on Whatmann paper and the parasite seeds were obtained by filtration through 50 and 100 mesh. Genomic DNA was extracted by PowerSoil kit (MO BIO Lab. Inc., Loker Ave West, Carlsbad, CA). according to the manufacturer’s instructions (Sagova-Mareckova et al., 2008).

### Quantitative real time PCR of soil sample from the sunflower fields

The quantitative real time PCR was performed using PerfeCTa® SYBR® Green Fast Mix®, ROX^TM^ (Quanta biosciences) and ABI-Prism 7000 Real-Time PCR Detection System (Applied Biosystems) according to the manufacturer’s protocol. The qPCR was performed in quantitative reaction with final volume of 10 µl including: 100ng DNA, 5 µl of SYBR Green Fast Mix-ROX and 500nM of each ITS-100 primers. *O. cumana* actin was used as endogenous control and the relative gene expression level was calculated using the 2^−ΔΔCt^ method (Livak and Schmittgen, 2001). Two standard curves were prepared: The first curve included standard points from *O. cumana* seeds ranging from 0.001 to 10 ng DNA per tube, were made using 1:10-fold serial dilutions of broomrape DNA and were used for qPCR assay. The second curve was generated by mixing different amounts of genomic DNA from *O. cumana* (0.1 to 100 ng) with total genomic DNA (10 µg) extracted from 250 mg soil sample. Each point on the standard curve was assayed in triplicate.

### Viability evaluation of the parasite seeds by Tetrazolium test

Tetrazolium test was performed as described earlier (Lopez-Granados and Garcia-Torres, 1999; Thorogood et al., 2009). Briefly, 50-100 seeds were hydrated between paper towel sheets for 2-3 days. After pre-conditioning, they were placed in flasks, covered with a 1% solution of 2,3, 5 triphenyl tetrazolium chloride (TZ) and incubated at 40°C for 2h. After that, the TZ solution was discarded; and the seeds were washed thoroughly with sterile water. Seeds were directly evaluated for viability under the microscope using magnification (40X and 100X) and classified in two categories: viable seeds – orange to brown light coloured or non-viable seeds –yellow to white colour.

## RESULTS

### Design of *rbcL* specific marker to differentiate between *O. cumana* and *O. crenata*

In our previous study, we reported amplification of ITS region using ITS350 primer; which specifically detect broomrape in soil sample (Aly et al., 2012). However, ITS350 marker does not differentiate between broomrape species because of the high sequence similarity among ITS regions. Therefore, we compared and analysed the *rbcL* gene sequences from these species using Clustal Omega and designed new molecular marker to differentiate broomrape species such as *O. cumana* and *O. crenata* (Fig. S1). To distinguish between the above broomrape species, probes differing at least by five-base mismatches or more were designed. Based on sequence alignment, a molecular marker (RbcL-M) was designed using *rbcL* gene from *O. cumana* (GenBank accession number: AF090349.1) Soil samples were artificially contaminated with seeds or shoots of *P. aegyptiaca, O. crenata* and *O. cumana* and subjected to PCR analysis. By PCR amplification using RbcL-M, we were able to differentiate between *O. crenata* and *O. cumana* according to the length of the PCR products. For *O. crenata* the PCR product size was 1300bp compared to 1000bp for *O. cumana*. However, RbcL-M marker failed to amplify any PCR product while soil samples were artificially contaminated with *P. aegyptiaca* seeds or shoots (Fig. 1a). As expected, the universal internal control primers amplified a PCR product (555bp) from soil samples with or without broomrape shoots or seeds (Fig. 1b).

**Fig. 1.**
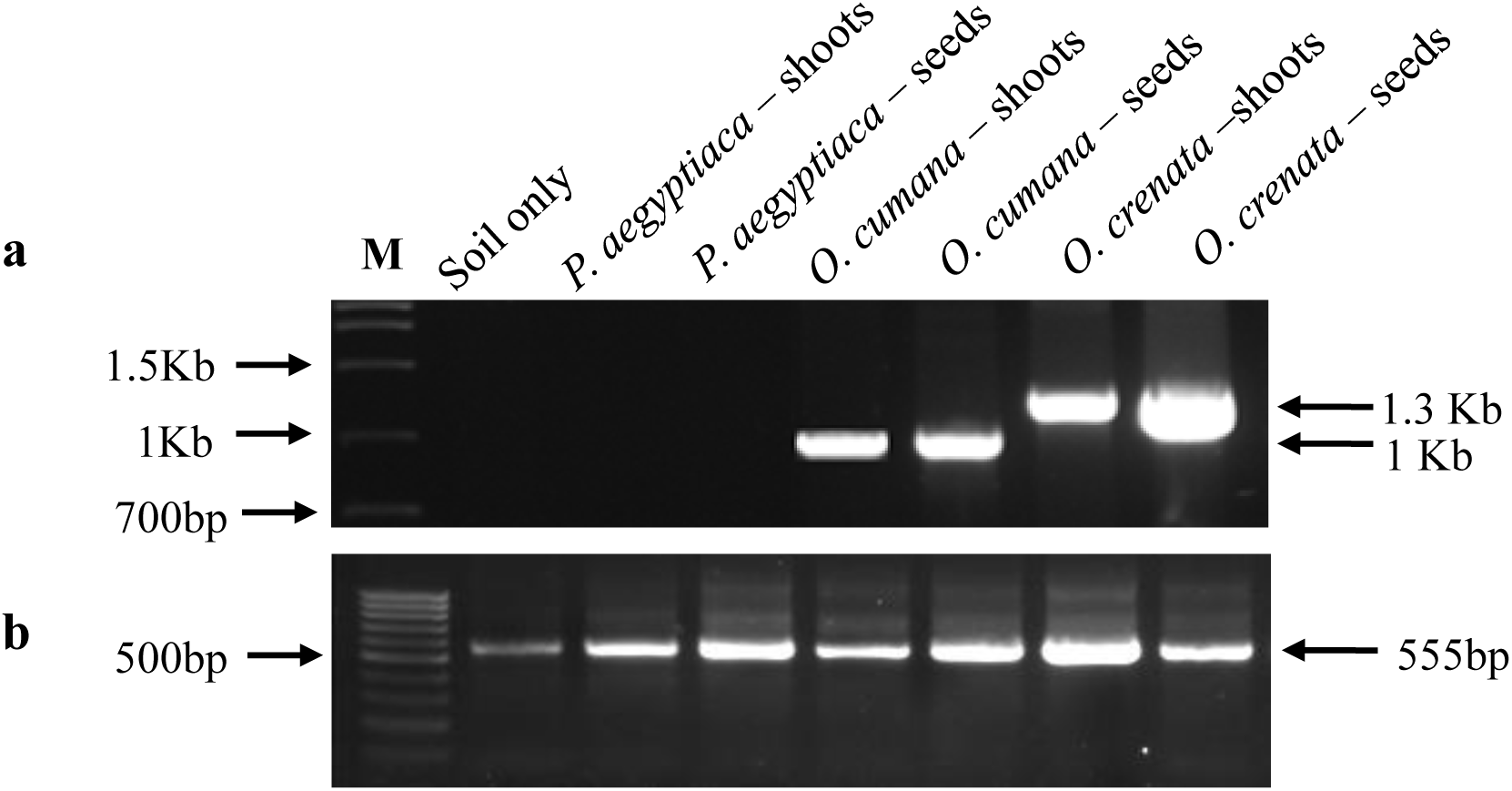
**(a)** Specificity of RbcL-M marker and PCR amplification patterns in three broomrape species (*P.aegyptiaca, O.cumana and O.crenata*). Soil samples (250 mg) were artificially contaminated with broomrape seeds (100) or 10 mg shoots of *P.aegyptiaca, O.crenata and O.cumana* and subjected to PCR analysis using RbcL-M primers. Genomic DNA was extracted and 100 ng were used for PCR amplification. A PCR product of 1300bp was obtained from *O.crenata* seeds and shoots, A PCR product of 1000bp was obtained from *O.cumana* seeds and shoots. However, RbcL-M marker failed to amplify PCR product with soil sample alone or with soil contaminated with *P.aegyptiaca* seeds or shoots. **(b)** The same samples were subjected to PCR detection using universal internal control primers (UCP-555). Arrows indicates the PCR product sizes.

### Development of specific ITS100 marker to detect seeds of three broomrape species in a soil sample

To analyse the number of the broomrape seeds present in a soil sample from infected field, we designed ITS100 primers from the highly conserved region of ITS350 region (Aly et al., 2012). To determine the specificity of the marker to each broomrape specie, we first extracted genomic DNA from soil samples (250mg) containing (0, 1, 10, 25, 50 100 and 250 seeds) of *P. aegyptiaca, O. crenata* or *O. cumana* and subjected to PCR analysis. ITS100 primers successfully amplified a PCR product (100bp) when *P. aegyptiaca, O. crenata* or *O. cumana* were mixed with soil sample, except soil sample without *Orobanche* seeds. However, a PCR product (555bp) was amplified in all samples including soil with or without broomrape seeds by using universal control primer UCP-555 (Fig. 2a).

**Fig. 2.**
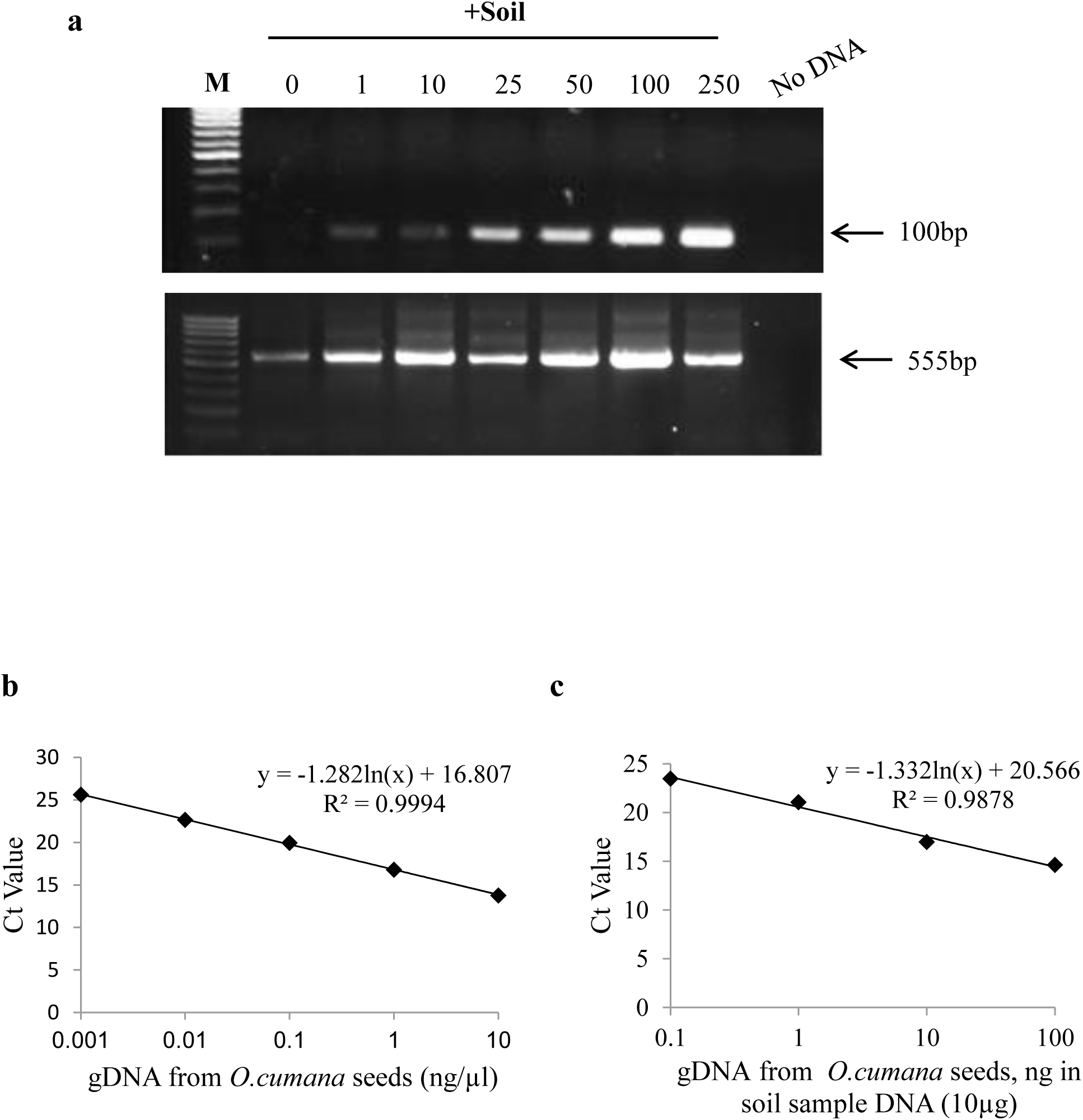
PCR amplification patterns of ITS100 primers in *O.cumana* seeds **(a)** gDNA was extracted from soil samples (250 mg) containing the parasite seeds (1, 10, 25, 50, 100 and 250 seeds). For each sample, 100 ng gDNA were used for PCR amplification. The same samples were subjected to PCR detection using universal internal control primers (UCP-555). Arrows indicates the PCR product sizes **(b)** Standard curves generated for qPCR to quantify *O.cumana* seeds in a soil sample. standard curve generated by plotting the value of threshold cycle value (Ct) against log of the amount of template DNA (ng/µl) from *O.cumana.***(c)** Detection and quantification of *O. cumana* template DNA (ng) from known amount of the parasite seeds mixed with extracted soil sample DNA (10 µg). Each point on the standard curve was assayed in triplicate.

In addition, the specificity of the markers generated in this study were evaluated by testing DNA extracted from sunflower, carrot and tomato leaves and roots corresponding to possible respond to the host plants. Fortunately, the qPCR assay always yielded negative results for these plant species (data not shown). Next to quantify *Orobanche*, we generated standard curve using *O. cumana* genomic DNA and ITS100 marker utilizing qPCR procedure. *O. cumana* genomic DNA was purified and serial dilutions (1:10 fold) ranging from 0.001 to 10 ng/µl were used for qPCR assay. The standard curve for *O. cumana* exhibited a slope of -1.28 and (R^2^) of 0.9994 (Fig. 2b). The same type of assay was performed with *O. cumana* genomic DNA extracted from soil sample (250 mg) mixed with the parasite seeds (0.1, 1, 10 and 100 mg seeds) and subjected to qPCR. The standard curve for *O. cumana* seeds mixed with soil sample, exhibited a slope of -1.332 and (R^2^) of 0.987 (Fig. 2c).

### Prediction and quantification of *O. cumana* seeds in sunflower field

Previously, we have developed a protocol that allows extraction of genomic DNA from few tiny seeds of *Orobanche* spp. in a soil sample and we were able to subject the sample DNA to a rapid PCR-assay (Aly et al., 2012). Here, we extended the protocol, allowing for the identification and distinguishing between soil-borne and parasite seed species that are collected from a field soil sample. Following confirmation and validation of the ITS100 marker specificity to *O. cumana* seeds, geo-statistics model for soil sampling that characterized the spatial variation in the field, was performed according to (Eizenberg et al., 2012) in a sunflower field located at Havat Eden, Emic Betshan– Israel, which was infected with *O. cumana* (Fig. 3a and b). Five soil samples (spot number 2,4,6,8 and 10) were collected from a plot of 1 acre according to the illustration (Fig. 3c). Genomic DNA was extracted from each soil sample and subjected to qPCR using ITS100 primers to predict and quantify *O. cumana* DNA in naturally infected field. Then, genomic DNA from soil samples was extrapolated with Ct values to quantify the number of seeds in each sample according to the generated Ct values (Fig. 2b and c). The results showed variable quantity of *O. cumana* seeds in the soil sample ranged from 2 to 22 parasite seeds (Fig. 3d) in 250mg soil sample processed originally from 1kg soil sample from the infected field. Sample no. 6 showed the highest density of *O. cumana* seeds (22 seeds) in the sunflower infected field as compared to the other samples. Sample no. 6 was collected from a highly infected area as was shown by the geo-statistics model proposed by (Eizenberg et al., 2012). Concomitantly, we monitored and counted the parasite inflorescences in adjacent to the position were the soil samples were taken. Our results revealed that average density of 0.6 inflorescences /m^2^, in a total area of 1acre was recorded. Inflorescences counts in the selected area in the field were also variable ranged from 2 to 15 and were less than the seed numbers found in the soil sample when qPCR was applied (Fig. 3d).

**Fig. 3.**
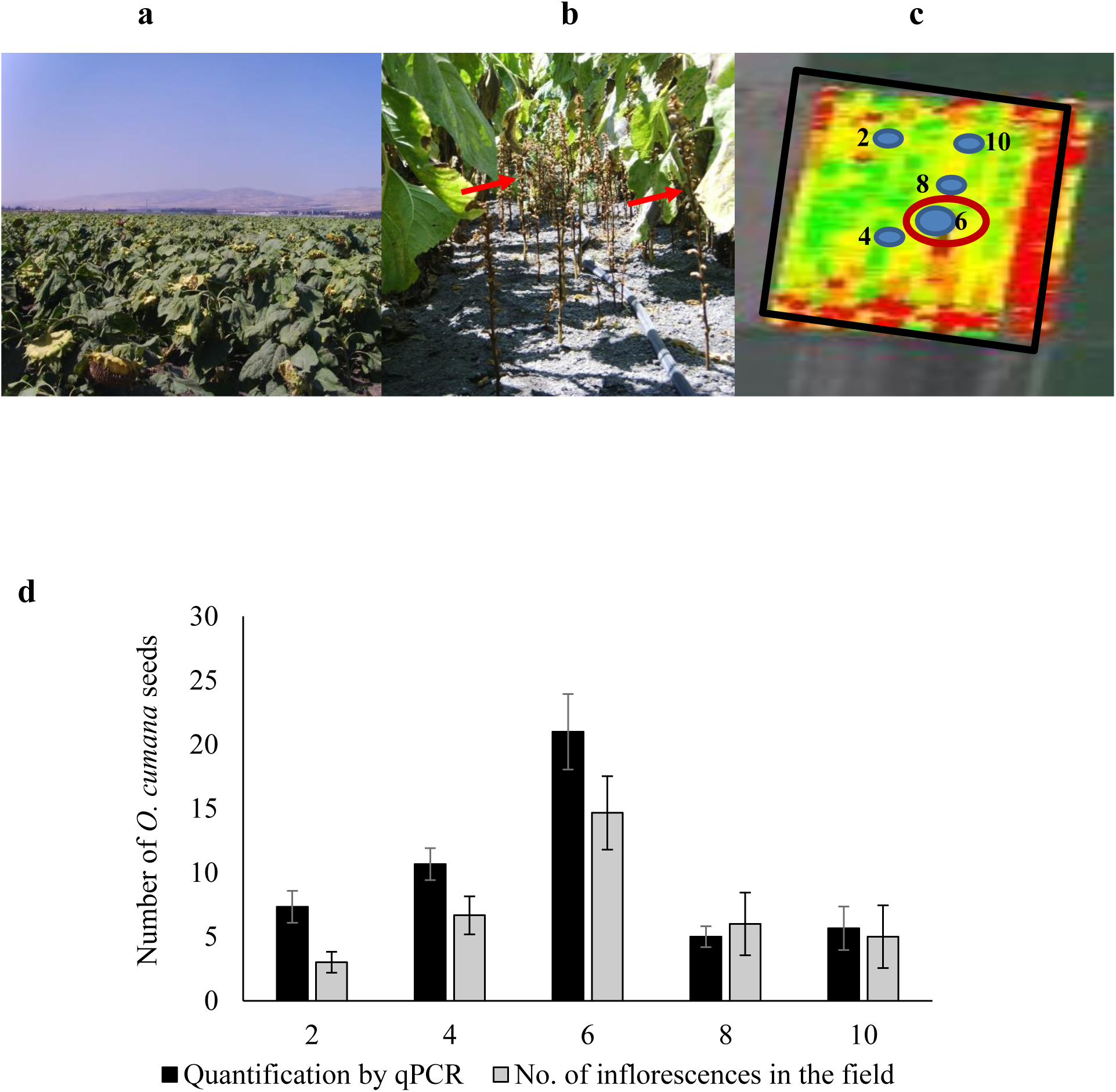
Quantification of *O.cumana* seeds in an infected sunflower field by qPCR and manually counting the inflorescences adjacent to the soil sample. **(a and b)** represents a section of the sunflower infected field. Red arrows show *O.cumana* inflorescences. **(c)** Representing a (GIS) model (Eizenberg et al., 2012) for soil sampling and characterize the spatial variation in sunflower field infected with *O.cumana* in Havat Eden. Five soil samples (2, 4, 6, 8, 10) of 500gm were collected from the infected field on May. Each sample was collected from a depth of 0 to 20 cm in a total area of 1×1 meter. The blue spots indicate location of the sample in the infected field. Red color in the plot represents high density of the parasite inflorescences. Infectivity was also monitored by counting the inflorescences on July and was restricted to the total area 1×1 meter of the five selected soil samples. **(d)** quantification of *O. cumana* seeds in a soil sample by qPCR (black bars) and monitoring the parasite inflorescences counts in fields (Grey bars). The data are the means of three soil samples or biological replicates. Vertical lines indicate SD of three independent measurements.

### Evaluation viability of the parasite seeds in sunflower infected field

To determine viability of broomrape seeds in a sunflower infected field, a tetrazolium test (TZ) (Lopez-Granados and Garcia-Torres, 1999; Thorogood et al., 2009) was conducted. According to TZ test, viable parasite seeds tends to have orange to brown colour because of its metabolite activity, while non-viable tends to have yellow to faint colour (Fig. 4a). To evaluate the parasite seed viability, two treatments were assayed: soil samples (organic matter only) collected from sunflower field located at Havat Eden, Emic Betshan, naturally infested with *O. cumana* and pure *O. cumana* seeds (1 mg) prepared from an old seed stock. Our results indicate that the highest percentage (% of total) of viable *O. cumana* seeds were found in soil samples collected from the sunflower field (83%) as compared to 25% found in the old seed stock (Fig. 4b).

**Fig. 4.**
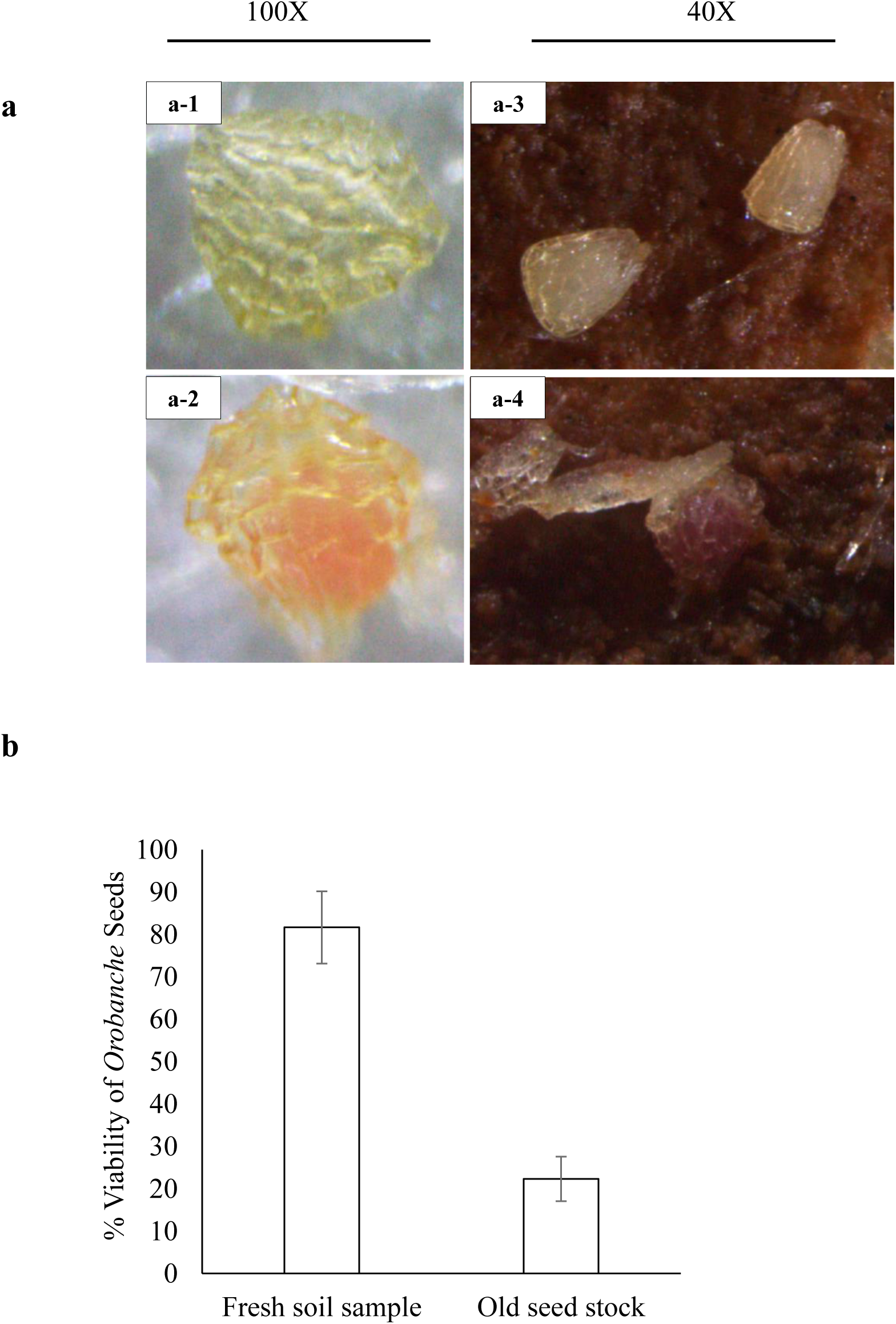
Evaluation the viability of *O.cumana* seeds by Tetrazolium test. **(a)** Tetrazolium test was used to visually differentiate between metabolically active tissue from (cleaned non-viable seed (a-1), non-viable seed with organic matter from an infected sunflower field (a-3), and active tissues from (cleaned viable seed (a-2), viable seed with organic mater from an infected sunflower field (a-4). Images were acquired using fluorescence microscope using 40X and 100X magnification. **(b)** The same test was also performed to evaluate percentage of viability of *O.cumana* seeds in soil samples from the infected field (83%) as compared to viability of the seeds from cleaned old seed bank (25%). The data are the means of three separate experiments with vertical lines indicating SD.

## DISCUSSION

Parasitic weeds are often exert their greatest damage prior to their emergence; therefore, the majority of field loss may occur before diagnosis of infection. Early detection of infection by visual inspection is not possible. This hampers early control strategy efforts and the detection methods used by crop scientists are laborious and time consuming. Most of the methods developed to detect broomrape seeds in soil samples, need several steps such as seed purification (flotation, filtration and binocular microscope observation (Kroschel, 2001). Despite of the great advances in genome sequencing approaches that has already been achieved, our understanding towards genomic evolution of Orobancheaceae is still incomplete due to aberrant evolution of plastid genome of holoparasite (Park et al., 2007). Available DNA sequence data are insufficient to differentiate closely related species, such as the weedy parasites *O. cumana, O. crenata* and *P. aegyptiaca*.

Genetic or DNA based marker techniques such as ITS or *rbcL* are routinely being used in evolutionary, taxonomical, phylogenetic and diversity studies (Agarwal et al., 2008; Park et al., 2008; Schneeweiss et al., 2004). Diagnosis and early identification of the parasite species in the field by soil sampling is of great importance to farmers and it is particularly needed due to the host-parasite specific interaction, the difficulty to detect the tiny parasite seeds by conventional methods in contaminated fields and because seeds will only germinate and grow in the presence of a susceptible host. The gene encoding the large subunit of ribulose-1,5-bisphosphate carboxylase/oxygenase (Rubisco; *rbcL*) is retained as a pseudogene in *Orobanche* and *Phelipanche* (Wicke et al., 2013). In the present work, we designed sets of primers targeting the plastid *rbcL* gene sequence from *O. cumana* to differentiate between three broomrape species (*O. cumana, O. crenata* and *P. aegyptiaca*) the most common and harmful species in field crops in Israel. Another sets of primers were designed to target ITS sequences (Schneeweiss et al., 2004) to quantify the parasite seeds potentially present in a soil sample to help for seed bank assessment in the field. We found that RbcL-M primers (marker) were useful to distinguish between DNA from *O. cumana* and *O. crenata* producing different amplicon sizes (a specific PCR product about 1300bp for *O. crenata* and 1000bp for *O. cumana*). Subsequent BLASTn with previously published broomrape genomic data (Cusimano and Wicke, 2016) reconfirmed the differences between the two species. In contrast, DNA from *P. aegyptiaca* failed to produce PCR amplicon suggesting possibly that *Phelipanche* species completely lost *rbcL* gene (Delavault et al., 1995; Leebens-Mack and de Pamphilis, 2002).To allow differentiation between *P. aegyptiaca* from other species, a specific molecular marker was recently designed (Aly et al., 2019), (GenBank accession numbers MK637618-637624).

To diagnose and quantify broomrape species in a seed stock (Dongo et al., 2012), different types of nuclear and plastid DNA markers have been proposed. (Schneeweiss et al., 2004) were the first to present molecular phylogenetic analysis using nuclear ITS sequences. ITS-based markers were also used to detect *P. aegyptiaca* seeds in a soil sample (Aly et al., 2012) and quantify contamination of *O. ramosa* and *O. cumana* in crop seed lots (Dongo et al., 2012). Here we developed ITS100 marker that was based on primers consisting of unique sequences in the internal transcribed spacer (ITS) regions of the nuclear ribosomal DNA (nrDNA) of *O.crenata*. ITS-100 marker was used with qPCR assay to quantify *O. cumana* seeds in a soil sample from sunflower field located at Havat Eden, Emic Betshan – Israel.

We were able to detect 0.1 mg *O. cumana* seeds in 250 mg soil sample (wt/wt). A detection threshold of 0.1 mg broomrape seeds in 20g seed samples was previously reported (Dongo et al., 2012). For detection, mapping and quantifying *O. cumana* seeds in the field, we used geographical information systems (GIS) for soil sampling and other advanced technologies for parasitic weed mapping and field history data storage (Eizenberg et al., 2012) followed by qPCR assay. Soil samples (500gr) were collected from the sunflower field, organic material was extracted from each sample ending with 250mg then, genomic DNA was extracted and subjected to qPCR using ITS100 primers. This method allowed specifically detecting and quantifying the DNA of *O. cumana* in a total DNA extract from sunflower soil sample. Accordingly, the results of this assay can be also expressed as the number of parasite seeds per kilogram of soil following extrapolation with the standard curve was prepared. Our results indicate that samples collected from highly infected area (Fig. 3c) according to the geo-statistics model proposed by Eizenberg et al. (2012), were with agreement with our qPCR assay (sample no. 6 showed the highest density of *O. cumana* seeds (22 seeds) in the sunflower infected field as compared to the other samples. However, no correlation was found between qPCR assay (parasite seed number) compared to number of the parasite inflorescences collected from the same location in the sunflower infected field. An explanation for that may be related to the viability of *O. cumana* seeds, genomic DNA from a soil sample will contain viable and non-viable parasite seeds, add to that, we cannot exclude the presence of some related parasite seeds in the same sample therefore, we counted more parasite seeds using the qPCR assay. Specificity of the qPCR assay was tested against several possible contaminants of soil-borne pathogens by using universal internal control primer UCP-555 (White, 1990) or harvested crop seeds like sunflower and tomato using ITS100. No amplification was observed, confirming the specificity of the marker.

The powerful of broomrape-infected field to distribute and contaminate the neighbouring non-infested fields depends on the soil seed-bank and viability of the parasite seeds. Viability of broomrape seeds in the sunflower infected field was determined by tetrazolium test. Our results showed the highest percentage of viable broomrape seeds were found in soil samples collected from the sunflower field (83%) as compared to 25% found in an old seed stock. We assume that the highest count recorded in the sunflower field was due to the release and distribution of the fresh parasite seeds by the newly parasite inflorescences showed up through the crop growth. We have to take in consideration that we used an old seed stock (10 years old) that doesn’t represent newly harvested seed stocks. Our experience with seed germination of *O. cumana* with germination stimulant (GR) could reach more than 90%. Additionally, we cannot exclude the presence of some related parasite seeds in the same sample from previous growth seasons.

In this study, we provide a simple, fast and non-expensive approach to distinguish and quantify broomrape seeds exist in a soil sample from a crop field. Molecular markers would assist accurate detection and population level of *Phelipanche* and *Orobanche* spp. in a soil sample and offer numerous advantages over conventional phenotype based alternatives, as they are stable and detectable in all tissues regardless of growth, differentiation or development. These methods could be helpful in precision agriculture, in which they provide answers routinely questioned by the farmers: are there parasite seeds in my crop field? What species? and how much seeds are exist in a soil sample?

## ACKNOWLEDGMENTS

Results of this research were supported by the Chief Scientist of the Ministry of Agriculture and Rural Development - Israel, grant No.132-1499-10 (MEZAMALEKET). VKB is grateful to the ARO-Volcani Center, Agricultural Ministry of Israel for providing the Postdoctoral fellowship.

## Supplementary Information

**Fig. S1.**
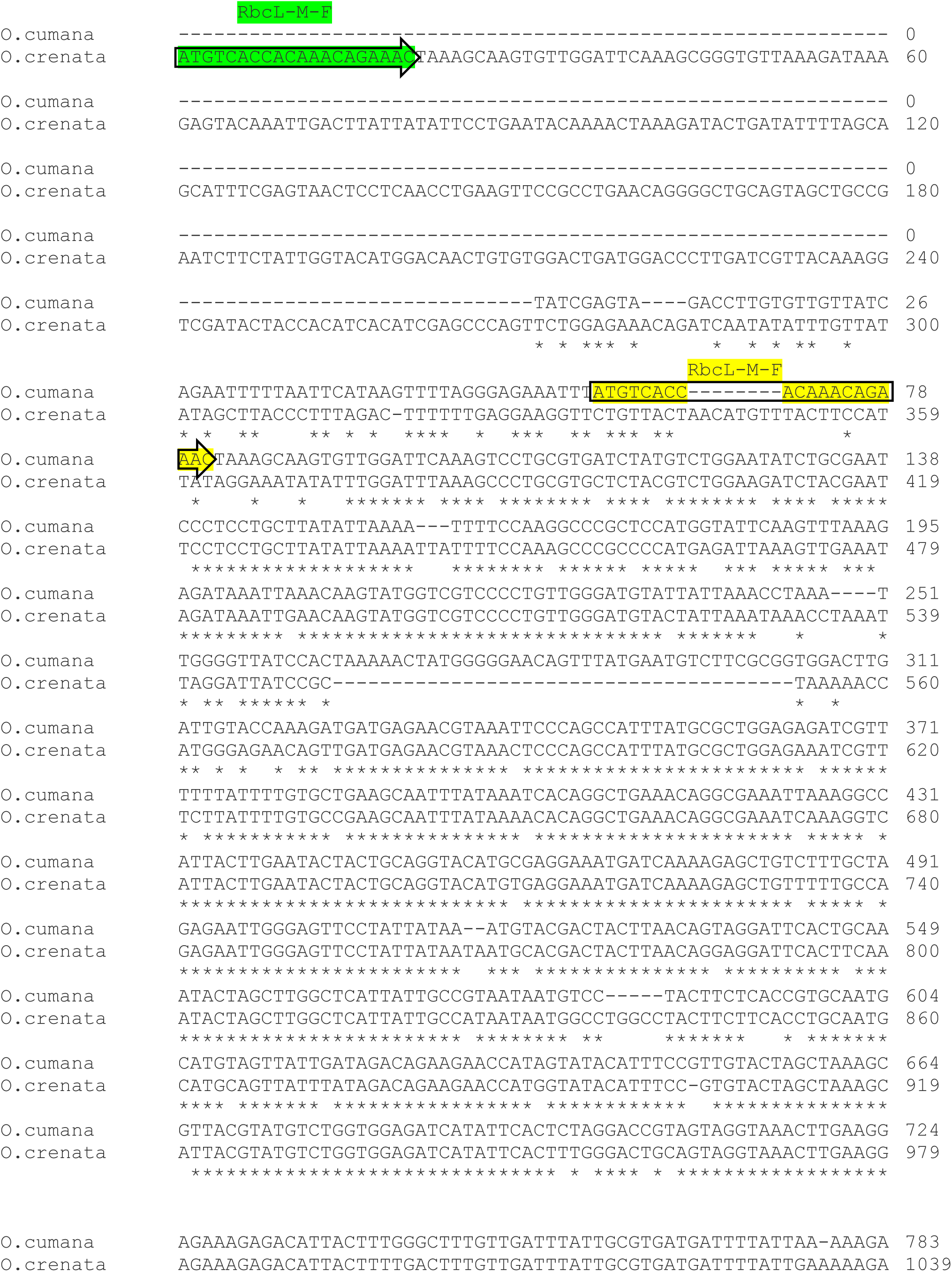
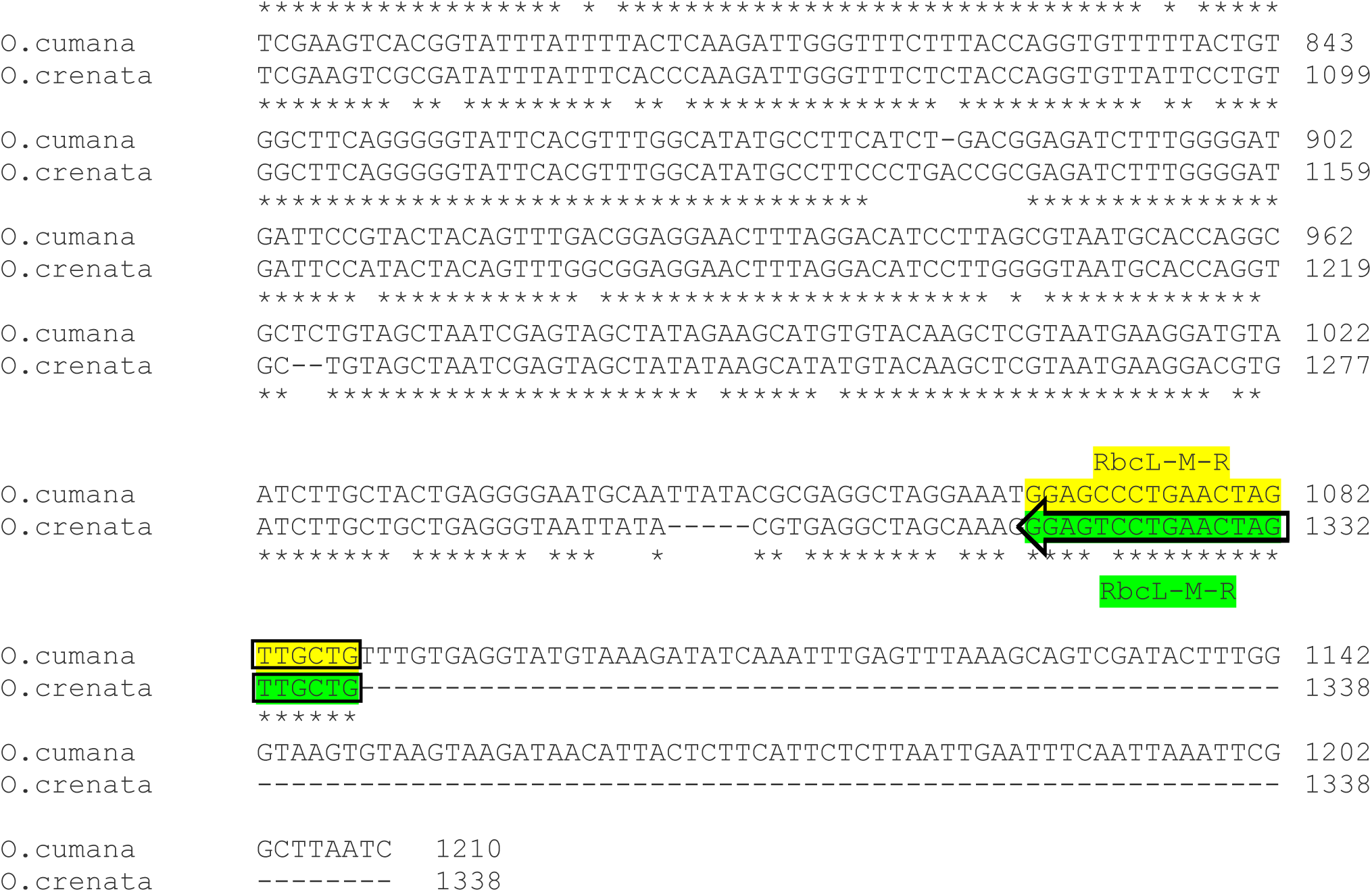
Multiple sequence alignment of *O.cumana* and *O.crenata rbcL* gene using online tool CLUSTAL Omega

**Table S1.**
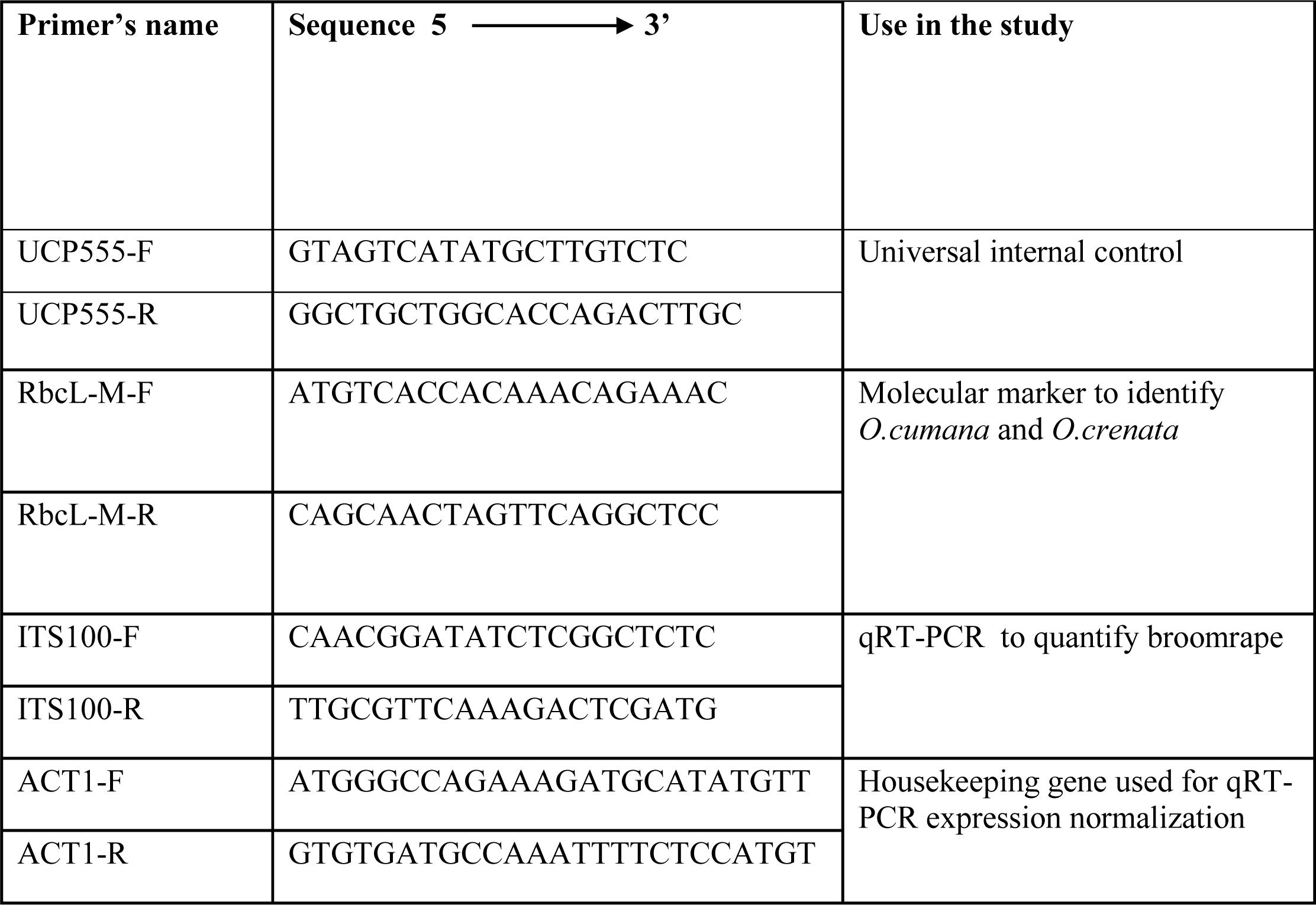
List of oligo’s used in this study

